# PELICAN: a Longitudinal Image Processing Pipeline for Analyzing Structural Magnetic Resonance Images in Aging and Neurodegenerative Disease Populations

**DOI:** 10.1101/2025.09.20.677546

**Authors:** Mahsa Dadar, Roqaie Moqadam, Amelie Metz, Katherine Chadwick, Aliza Brzezinski-Rittner, Yashar Zeighami

## Abstract

Structural magnetic resonance imaging (MRI) allows for accurate non-invasive assessment of the brain’s structure and its longitudinal changes. Availability of large scale longitudinal MRI datasets enables us to probe brain changes in health and disease, and derive longitudinal trajectories of brain morphometry based measures to estimate brain atrophy and other disease-related abnormalities. In contrast to their cross-sectional counterparts, image processing pipelines that have been designed for longitudinal data can reduce noise in the derived measurements by disentangling the within and between subject variabilities, improving the sensitivity of the downstream models in detecting more subtle longitudinal changes. Here we present PELICAN, our open source multi-contrast longitudinal image processing pipeline, that has been designed and extensively validated for use in longitudinal settings and populations with neurodegenerative disorders. PELICAN can use population specific average templates as intermediate targets to derive accurate nonlinear registrations for cases with substantial levels of atrophy, which commonly used pipelines struggle to process. We evaluated PELICAN’s performance across over 34,000 MRIs from multiple aging and neurodegenerative disorder cohorts, and compared its reliability and failure rates against FreeSurfer as a widely used image processing tool, showing superior performance of PELICAN compared to FreeSurfer, both in terms of failure rate and reliability. Our results demonstrate that PELICAN can be used to accurately process MRIs of individuals with neurodegenerative disease who present with greater levels of atrophy and white matter lesion burden.

## Introduction

Longitudinal magnetic resonance imaging (MRI) studies of aging and neurodegenerative disease populations can provide invaluable information on the severity, pattern, and evolution of pathology characteristic of different disorders such as Alzheimer’s disease^1–3^, Parkinson’s disease^4,5^, frontotemporal dementia^1,6^, Amyotrophic Lateral Sclerosis (ALS)^7^, and Multiple Sclerosis (MS)^8^. Open access and freely available image analysis pipelines allow for detailed assessments of the brain, and facilitate study replications and reproducibility^9–11^. While processing each individual time point independently can provide useful morphometry information and might be suitable in certain experimental designs (e.g. in predictive analyses where time points should be assessed independently to avoid leakage^12^), using longitudinal image processing pipelines can reduce measurement noise and provide more accurate downstream estimates^13^.

Another important concern when analyzing data from disease populations is the presence of pathology and how that might impact the derived estimates. Tools that have been designed and validated solely based on homogeneous populations (e.g. healthy young adults) might be less accurate when applied to diseased populations where severe pathology is present. For example, white matter hyperintensities (WMHs) are important MRI-visible pathologies, commonly present in aging individuals, and more so in patients with neurodegenerative disorders^14,15^. While they are more visible on T2-weighted (T2w) and fluid attenuated inversion recovery (FLAIR) images, they appear as hypointensities on T1-weighted (T1w) MRIs, and can have intensity ranges that are similar to cortical and deep gray matter^15–17^. We and others have shown that image analysis pipelines that have not been specifically designed and validated to handle the presence of WMHs can segment them as gray matter^18^, and that these errors propagate to all downstream analyses and can systematically bias study findings as WMHs are more prevalent in many diseased populations^17^.

Many image processing pipelines compare and standardize the individual MRIs to a standard template^11,13^ or use it to represent their results in a common space for further statistical analyses. Registration to a template is an inherent step in deformation-based (also referred to as tensor-based morphometry)^19,20^ and voxel-based morphometry analyses^21,22^. The most commonly used standard template is the MNI-ICBM2009, which is an average of 152 predominantly male young healthy adults^23,24^. However, use of young healthy templates for analyzing aging and diseased populations can lead to significantly greater linear and nonlinear registration errors^9,25,26^. These errors are generally most pronounced in cases with more severe pathology where the individual image contains significant neuroanatomical differences compared to the template, leading to inevitable exclusion of the more advanced disease cases after quality control and biasing the downstream analyses towards the healthier individuals. On the other hand, while using population-based templates specific to each study provides an unbiased estimate of atrophy, it also makes it challenging to compare results across studies (e.g., meta-analysis studies) without re-analyzing the data. Further still, calculating population-based templates for large scale datasets with thousands of participants can become computationally prohibitive.

In this work, we present our <u>P</u>ipeline for <u>E</u>valuating <u>L</u>ongitudinal <u>I</u>mages of <u>C</u>erebral <u>AN</u>atomy (PELICAN), an open source longitudinal image processing pipeline that has been designed and validated to perform accurate image registration, segmentation^27^, and deformation-based and voxel-based morphometry for cross-sectional and longitudinal analyses of multi-modal MRIs in aging and neurodegenerative disease populations. We validate the performance of PELICAN against FreeSurfer^11^, one of the most commonly used open access brain imaging pipelines that provides both cross-sectional and longitudinal image processing.

## Methods

### Data

T1w, T2w, and FLAIR MRIs from several large open access multi-center and multi-scanner datasets were used to validate the performance of PELICAN, including the UK BioBank^27^, the National Alzheimer’s Coordinating Center (NACC)^28,29^, the Alzheimer’s Disease Neuroimaging Initiative (ADNI)^30,31^, and the frontotemporal lobar degeneration neuroimaging initiative (NIFD)^6^. These datasets were selected to allow us to assess the performance of PELICAN on data from different imaging protocols, age ranges, and neurodegenerative pathologies.

#### UK BioBank

Participants were included from the UK Biobank, an open-access large prospective study with phenotypic, genotypic, and neuroimaging data from 50,000 individuals recruited between 2006 and 2010 at 40–69 years old in Great Britain; data collection was performed across three different sites^32^. All participants provided informed consent. The UK Biobank received ethical approval from the Research Ethics Committee, and the present study was conducted based on application 45551.

#### ADNI

Participants were included from the ADNI database. The ADNI was launched in 2003 as a public-private partnership, led by Principal Investigator Michael W. Weiner, MD. The current dataset consists of over ∼16,000 MRI visits from ∼2,400 elderly controls, individuals with mild cognitive impairment (MCI), and Alzheimer’s dementia^31^. The primary goal of ADNI has been to test whether serial MRI, positron emission tomography (PET), other biological markers, and clinical and neuropsychological assessment can be combined to measure the progression of MCI and early AD. All participants provided informed consent and the study protocol was approved by the institutional review boards at all sites. A random subset of 388 images were included in this study.

#### NIFD

NIFD was founded through the National Institute of Aging and started in 2010^33^. The primary goals of NIFD were to identify neuroimaging modalities and methods of analysis for tracking frontotemporal lobar degeneration and to assess the diagnostic value of imaging versus other biomarkers. NIFD includes 199 participants with different variants of frontotemporal dementia and 136 healthy controls, recruited at three research sites across the United States. All participants provided informed consent and the protocol was approved by the institutional review boards at all sites.

#### NACC

We also included T1w, T2w, and FLAIR MRI data provided by the National Alzheimer’s Coordinating Center’s Uniform Data Set (NACC)^28,29^. NACC has aggregated a database of standardized clinical research data obtained from past and present NIA-funded Alzheimer’s Disease Research Centers across the United States, allowing us to assess the performance of PELICAN in non-harmonized datasets with a wide range of scanning resolutions and protocols.

#### SIMON

To assess test-retest reliability of PELICAN, 57 T1w MRIs that were acquired within a one year interval (age 45-46) were included from the SIMON dataset, a convenience sample of one healthy male aged between 39 and 46 years old, scanned for research projects in 73 sessions at 28 sites on a variety of scanner manufacturers and models: GE Healthcare (DISCOVERY MR750 and SIGNA Pioneer); Philips Medical Systems (Achieva, Ingenia, Intera, and T5); and Siemens Healthcare (Allegra, Prisma, PrismaFit, Skyra, SonataVision, Symphony, and TrioTim)^34,35^. Ethics approvals were obtained at all respective sites and the participant provided written informed consent to release his data without restriction.

Table 1 summarizes the characteristics of the participants and MRIs included from each dataset.

**Table 1.** Characteristics of the participants and MRIs included in this study.

| Dataset | UK BioBank | NACC | NIFD | ADNI | SIMON |
| --- | --- | --- | --- | --- | --- |
| N (N<sub>Female</sub>) | 24,732 (13355) | 2,951 (1650) | 329 (154) | 388 (248) | 1 (0) |
| N<sub>Time points(per subject)</sub> | 28,740 (1 - 2) | 4,446 (1 - 4) | 751 (1 - 4) | 843 (1 - 6) | 53 (53) |
| Age ($\mu \pm \sigma$) | 64.0 ± 7.67 | 70.98 ± 10.88 | 63.50 ± 7.29 | 73.21 ± 7.03 | 45.6 ± 0.6 |
| Min - Max | 44-82 | 18 -100 | 39 - 86 | 55 - 91 | 45-46 |
| Diagnosis | Aging | Aging, MCI, AD | Aging, FTD | Aging, MCI, AD | HC |
| T1w Resolution | 1.0 × 1.0 × 1.0 | 0.8-1.2 × 0.8-1.2<br>× 0.8-1.2 | 1.0-1.4 × 1.0-1.4<br>× 1.0-1.4 | 1-1.2 × 0.9-1.25<br>× 0.9-1.25 | 1.0 × 1.0 × 1.0 |
| T2w Resolution | - | - | - | 3 × 0.93 × 0.93 | - |
| FLAIR Resolution | 1.0 × 1.0 × 1.0 | 1-3 × 0.5-1.2<br>× 0.5-1.2 | 0.5-1.0 × 0.5-1.0 ×<br>0.5-1.0 | 5 × 0.85 × 0.85 | - |

T1w: T1-weighted. T2w: T2-weighted. FLAIR: fluid attenuated inversion recovery. PD: Parkinson’s Disease. AD: Alzheimer’s Dementia. MCI: Mild Cognitive Impairment. FTD: Frontotemporal Dementia. M: mean. $\sigma$ : Standard deviation.

### PELICAN

PELICAN has been developed based on the open access MINC Toolkit-v2 (http://bic-mni.github.io/) and ANTs (https://antsx.github.io/ANTs/) toolkits and can take either.mnc or.nii images as inputs. The pipeline is designed to process both cross-sectional and longitudinal T1w and any combination of T2w, Proton Density, and FLAIR MRIs; i.e. inclusion of T1w images is necessary for all time points, whereas other modalities are optional. Figure 1 shows the workflow of the pipeline steps, starting with preprocessing and co-registration of the available modalities, followed by linear registration to an average template, brain extraction, nonlinear registration to the average template, tissue segmentation, deformation-based and voxel-based morphometry, and creation of quality control images for all steps. In case of longitudinal data, the pipeline also includes construction of linear and nonlinear subject specific templates and additional steps of registration between the subject specific templates to the average template, providing both direct and indirect registrations for all time points.

**Figure 1.**
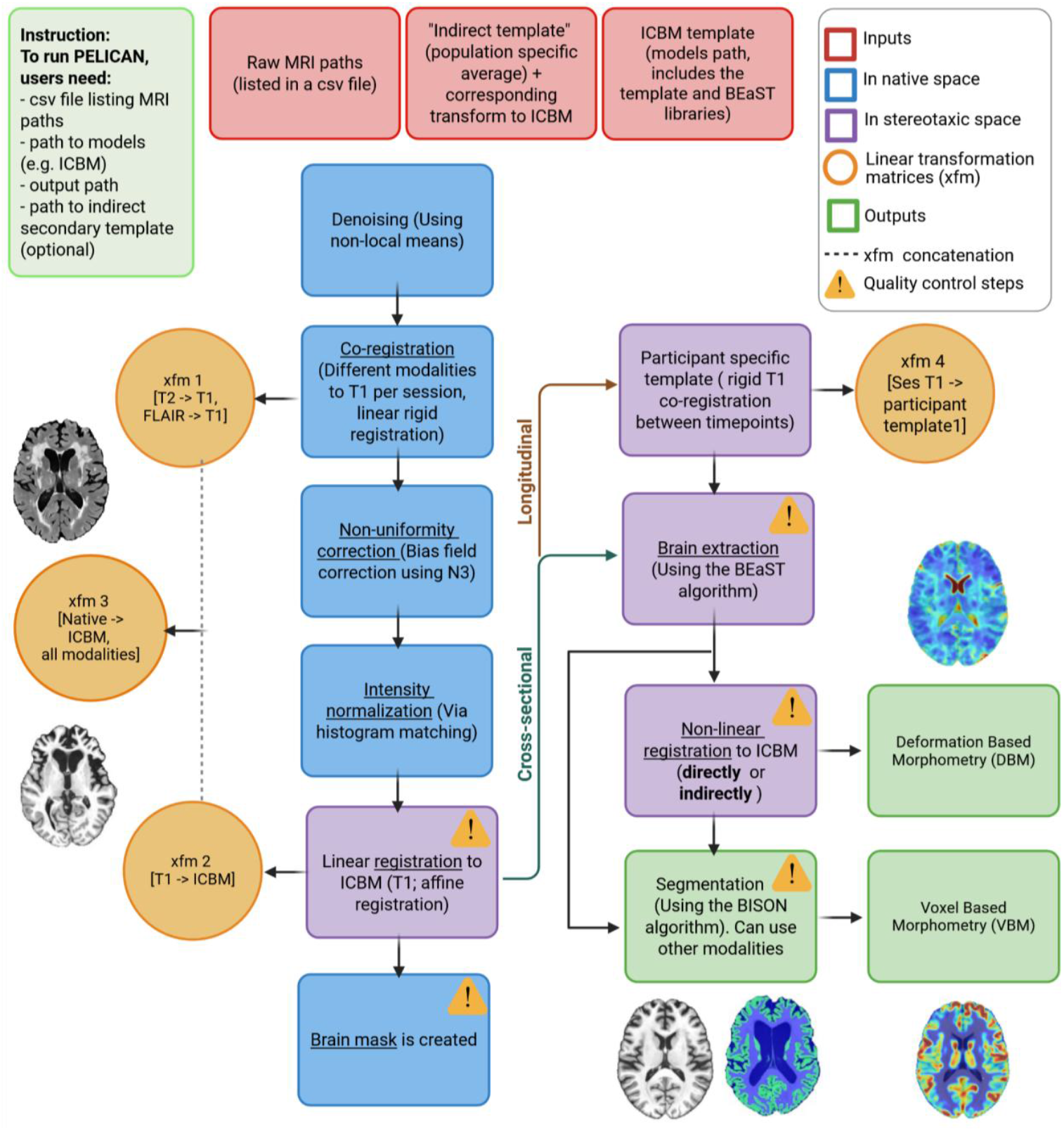
Workflow of PELICAN. The pipeline performs preprocessing, followed by linear registration, brain extraction, nonlinear registration, tissue segmentation, and deformation and voxel based morphometry analyses and generates quality control images for each step.

### Preprocessing

All available modalities are first denoised in their native spaces using the non-local means denoising *mincanlm* tool^36^. The next preprocessing steps are image intensity non-uniformity correction and intensity normalization, which are also best performed in the native image space and using a co-registered brain mask. To generate this native mask for all modalities, each modality is first co-registered to their corresponding T1w image from the same time point using *antsRegistration_affine_SyN* linear rigid registration^10^ (i.e., using 6 parameters). The T1w image is also linearly registered to the provided average template using an affine *antsRegistration_affine_SyN* registration step. The two transforms are then concatenated to provide native-to-template linear transforms for all available modalities. Using these transforms, the brain mask of the average template is resampled to the native spaces of each modality to provide a corresponding native brain mask to be used for the non-uniformity correction step. Note that all the brain masks and transformations in this step are temporary, and will be recalculated after the preprocessing is completed to improve their accuracy. Intensity non-uniformity correction is performed on the denoised images using *nu_correct*, which is a MINC-based N3 implementation^37^. Intensity normalization is performed on the non-uniformity corrected images to bring them into an intensity range of 0-100, using the MINC-based *volume_pol* linear intensity histogram matching method with the same brain masks and corresponding modality-specific average templates as input (i.e. intensity ranges for each modality are normalized based on population average images of the same modality).

### Registration

Once all available modalities are preprocessed in their native spaces, the linear rigid co-registration steps are performed again on the preprocessed images to improve their accuracy. In case of cross-sectional data, the preprocessed T1w image is also registered to the average template again and the resulting transformations are concatenated and used to resample all available modalities to the stereotaxic space, to minimize resampling interpolation error. The MINC-based *BEaST* brain extraction tool^38^ is applied to the linearly registered T1w image to generate a brain mask to be used for nonlinear registration. Nonlinear registration to the population template is performed using the linearly registered T1w image and *BEaST* mask as inputs to the *antsRegistration* tool from ANTs^10^. In case of longitudinal data, the native-to-template registration steps utilize a subject-specific average template to improve their accuracy, as described in the following section.

### Subject-Specific Average Template Generation

Following preprocessing, the longitudinal T1w images of each participant are used to create an unbiased subject-specific T1w average template, similar to previous longitudinal image processing pipelines^13,22,39^. To achieve this, all time points are first linearly registered to the first time point, and a first-level linear average template is generated based on these images. Then, all images are linearly registered to the resulting average template and a new linear subject-specific template is generated based on the resulting registrations in 4 iterative steps. The linear subject-specific template is then registered linearly to the population average template, and the transformation is used alongside the timepoint-to-subject-average transformations in the last step of the linear template creation to bring all time points to the stereotaxic space. A similar hierarchical process is then repeated to generate a nonlinear subject-specific template in the stereotaxic space for each participant, using the linear subject-specific template as the initial step, and using the nonlinear subject-specific template generated in the previous step as the registration target for the next step afterwards. Nonlinear registrations are performed at a coarser level in the initial step, and refined in an iterative process, for a total of 4 steps (i.e. shrink-factors and smoothing-sigmas are reduced at each step), using *antsRegistration* tool from ANTs^10^. Transformations from each individual time point to the population template are calculated by concatenating the individual time point to the subject-specific template and the subject-specific template to the population appropriate average template^25,26^.

### Indirect Average Template Registration

While use of well-known templates such as the MNI-ICBM152 as average template has the advantage of facilitating interpretation of the results (e.g. reporting MNI coordinates corresponding to known anatomical landmarks for significant regions), allowing application of commonly used atlases for deriving regional measurements, and combining results across multiple analysis across cohorts, it can come at the expense of increasing registration errors, particularly during nonlinear registration and in populations where severe pathology is present^9,25,26^. Further still, experimental design might require combining data from different populations, each with their own characteristic pattern of atrophy that would hinder use of a specific disease population template. We have thus implemented an indirect average template registration step in our pipeline, that allows for use of a second population specific template as an intermediate registration target for the nonlinear registration step. If provided, nonlinear registration is also performed with this template as an intermediate registration target, and the final nonlinear transformation is calculated by concatenating the individual-to-indirect-template and indirect-template-to-average-template nonlinear transformations. Note that this step is performed in addition to the direct individual-to-template registrations and quality control images are provided for both approaches to allow the users to select the most appropriate outputs for their application of interest. We provide two indirect templates for Alzheimer’s disease and frontotemporal dementia populations with the pipeline which have proved useful for enhancing nonlinear registration quality based on our experience. We have further generated open access average templates for several other neurodegenerative disease populations^25,26^ that can be used as indirect templates for PELICAN. Furthermore, using new indirect-templates as inputs, PELICAN itself can be used to produce the necessary indirect-template-to -average-template transformations and segmentations for this additional step.

### Tissue Segmentation

The *BISON*^18^ tool is used to segment the processed images and generate tissue probability maps to be used to perform voxel-based morphometry. BISON is particularly suited for this task as it can provide reliable^35^ segmentations in development, young adult, aging, and neurodegenerative disease populations^18^. Furthermore, BISON can perform segmentations for both healthy tissue types and WMHs based both solely on T1w images and on multi-modal combinations of inputs (e.g. T1w+ T2w or T1w + FLAIR)^16,18^. We used a version of *BISON* that provides the following 9 labels as outputs: ventricles, cerebrospinal fluid (CSF), subcortical cerebral gray matter, cortical cerebral gray matter, white matter, WMHs, brainstem, cerebellar gray matter, cerebellar white matter. *BISON* outputs are merged into 3 tissue maps (gray matter, white matter, and CSF) for further use in calculating voxel-based morphometry maps.

### Deformation-Based and Voxel-Based Morphometry

Deformation-based morphometry (DBM) can be used to identify anatomical changes by spatially normalizing T1w MRIs to conform to the same stereotaxic space. DBM maps are calculated as the Jacobian determinant of the inverse deformation field^19^. Jacobian determinant values reflect the relative volume of the voxel to the average template; i.e. a value of 1 indicates similar volume to the template, values lower than one indicate volumes smaller than the corresponding region in the template, while values higher than one indicate volumes that are larger than the corresponding region in the template. Therefore, a decrease in the Jacobian determinant values for a specific region between two timepoints can be interpreted as reduction in the structure volume, i.e. atrophy. The nonlinear registrations calculated in previous steps and the MINC-based *grid_proc* tool are used to generate deformation-based morphometry (DBM) maps. When indirect templates are provided as intermediate registration targets, DBM maps are calculated and provided based on both the direct as well as the indirect nonlinear transformations to allow users to select the best outputs based on quality control results. The *BISON*-derived tissue probability maps and the DBM maps are used in combination with the MINC-based *mincblur* tool to derive voxel-based morphometry (VBM) maps.

### Quality Control Images

The MINC-based *minc_qc* tool is used to generate a series of quality control images for different steps of the pipeline. These include images showing axial, coronal, and sagittal slices of all available modalities linearly registered to the average template, with the contour of the template overlaid on them to allow for accurate assessment of the linear registration quality, similar to our previous work^9^. A quality control image is also generated for the *BEaST* brain mask by overlaying it on the linearly registered T1w images. Nonlinear registration quality control images are generated by applying the nonlinear transformations (both direct and indirect, if available) to each T1w time point, and overlaying the contour of the average template on the resulting image. *BISON* generates its own quality control images, which overlay the 9 tissue labels on the T1w image. Finally, to facilitate quality control of the outputs, all the quality control images are compiled into an html summary report along with description of each image and notes on what details should be verified in each step. Example images for each step are provided in the Supplementary Materials (Figures S.1 to S.10).

### Pipeline Validation

To validate the performance of the PELICAN and ensure its generalizability, we applied it to multi-center and multi-scanner data of over 24,000 aging and diseased individuals (over 35,000 time points) and performed visual quality control for all steps using the generated quality control images. All quality control steps were blind to participant diagnosis information. We tested the performance of the pipeline using both cross-sectional and longitudinal data, with and without using indirect templates. To assess whether the indirect template option of the pipeline improves the nonlinear registrations, quality control failure rates were compared for direct versus indirect nonlinear registrations in NIFD (using a frontotemporal dementia population template) and ADNI (using an Alzheimer’s disease population template) datasets, using chi-squared tests.

We further analyzed and quality controlled a subset of this data using FreeSurfer version 7.4.1, to provide comparisons against a commonly used image processing pipeline that has both cross-sectional and longitudinal options. *FreeSurfer* results were quality controlled using a consistent process and protocol, namely, the same quality control images were generated for *FreeSurfer* registrations and tissue segmentations. For the segmentations, *FreeSurfer* labels were merged to match the nine *PELICAN* labels to enable comparisons. Failure rates across different scenarios and datasets were compared using chi-squared tests.

### Reliability Assessments

To assess and compare the reliability of the estimated measurements, *PELICAN* and *FreeSurfer* were run cross-sectionally on 53 T1w images from the SIMON^34,35^ dataset that were acquired on different MRI scanners and with different protocols during a one year interval, and variability in the estimated intracranial volumes (eTIV), cortical and deep gray matter, white matter, and WMH volumes were compared between the two measurements.

### Data and Code Availability

All the datasets used can be directly requested through their respective online platforms. The code for the *PELICAN* pipeline as well as a singularity build for all its necessary tools is open source and publicly available at https://github.com/VANDAlab/Preprocessing_Pipeline.

## Results

Table 2 summarizes failure rates of the different major *PELICAN* steps for each dataset. Figure 2 provides examples of a case that passed quality control for all steps, and cases that failed quality control for each step. Failure rates were consistently below 4% for all steps and datasets, except for direct nonlinear registrations in NIFD (6.34%) and ADNI (5.88%), where extensive atrophy levels led to higher failure rates. However, using indirect nonlinear registrations to the MNI-FTD-136 and ADNI average templates as intermediate registration targets, PELICAN failure rates were significantly reduced to 1.83% and 1.86% (p < 0.05), respectively. Figure 3 compares a case that failed quality control for direct nonlinear registration to the MNI-ICBM-152 template, and passed quality control for indirect nonlinear registration using the MNI-FTD-136 template. Note that for NIFD, all raw images were QCed prior to running PELICAN, and the best available image was selected to be processed for each modality and visit, and cases that did not pass this QC step were not analyzed, whereas for the other datasets, this step was not performed. This might explain the lower failure rates for NIFD. It is also important to note that the failure rates for each step are reported based on cases that have passed QC for all prior steps. For example, nonlinear registration failure rates are reported based on cases that have passed linear registration and brain mask steps, since a failure in either of those steps would inevitably lead to inaccurate nonlinear registrations.

**Table 2.**
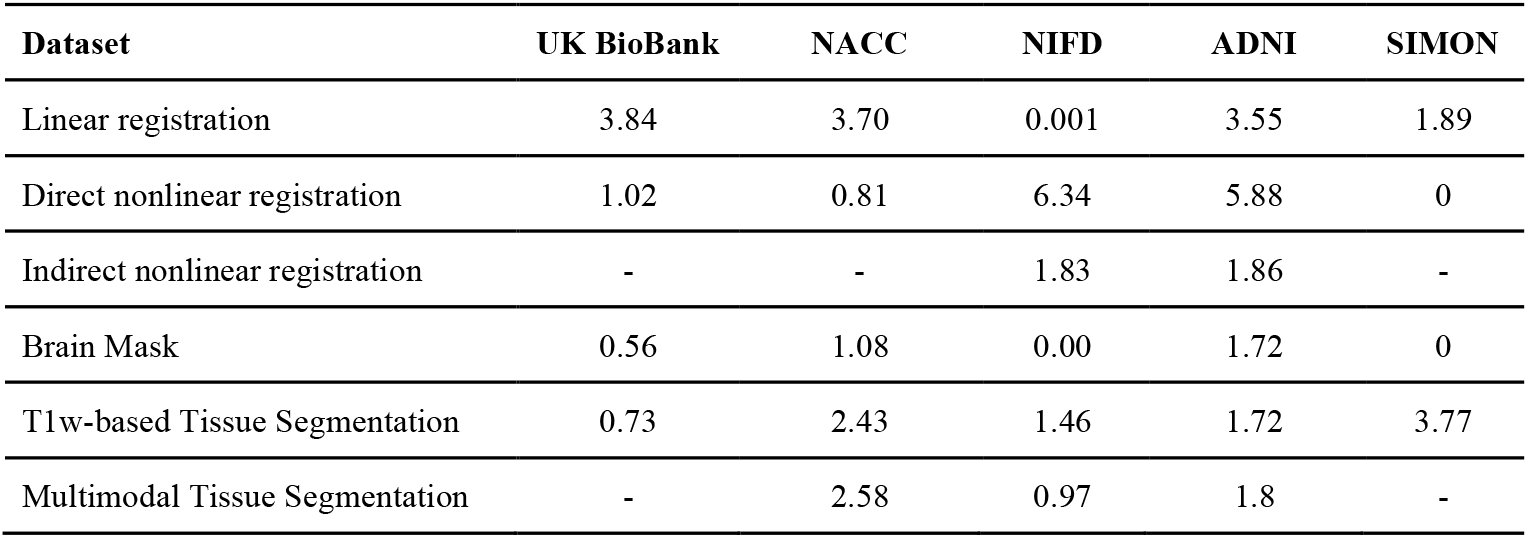
PELICAN failure rate percentages for the datasets included in this study. Note the higher failure rate of direct nonlinear registration in the NIFD and ADNI datasets, and how adding the indirect step mitigates this issue.

| Dataset | UK BioBank | NACC | NIFD | ADNI | SIMON |
| --- | --- | --- | --- | --- | --- |
| Linear registration | 3.84 | 3.70 | 0.001 | 3.55 | 1.89 |
| Direct nonlinear registration | 1.02 | 0.81 | 6.34 | 5.88 | 0 |
| Indirect nonlinear registration | - | - | 1.83 | 1.86 | - |
| Brain Mask | 0.56 | 1.08 | 0.00 | 1.72 | 0 |
| T1w-based Tissue Segmentation | 0.73 | 2.43 | 1.46 | 1.72 | 3.77 |
| Multimodal Tissue Segmentation | - | 2.58 | 0.97 | 1.8 | - |

**Figure 2.**
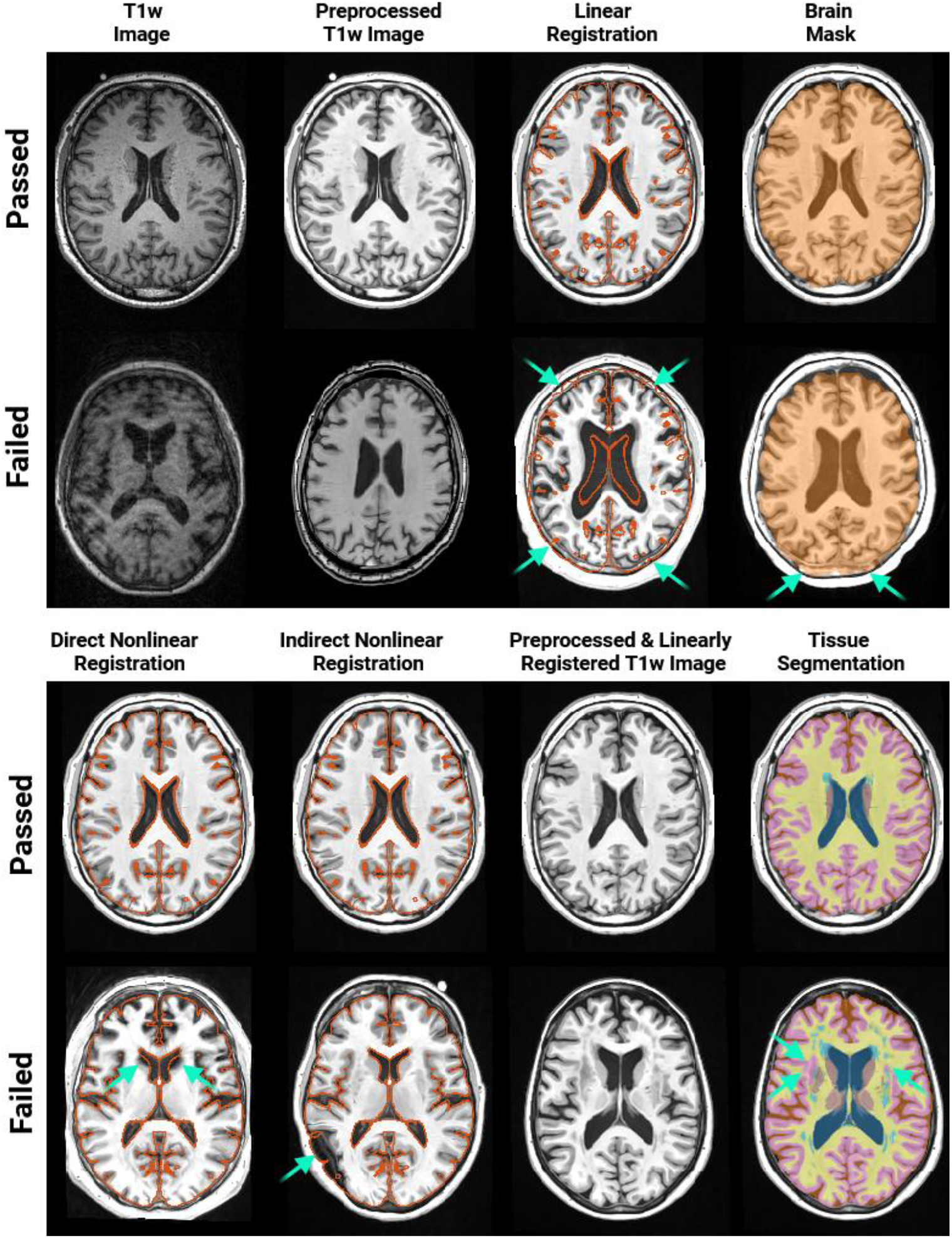
**Top rows:** example of a case that passed quality control for all PELICAN steps. **Bottom rows:** Examples of cases that failed each step. From left to right, causes of failure were: **T1w Image:** participant with motion during acquisition. **Preprocessed T1w:** failed intensity normalization. **Linear Registration:** inaccurate estimation of the scaling factor, note the alignment of the MNI-ICBM152 contour indicated in red. **Brain Mask:** inaccurate segmentation in the posterior regions. **Direct Nonlinear registration:** inaccurate registration in the anterior portions of the lateral ventricles. **Indirect Nonlinear Registration:** inaccurate registration in the left parietal lobe. **Tissue Segmentation:** inaccurate segmentation of WMH areas adjacent to the left putamen as cortical GM.

**Figure 3.**
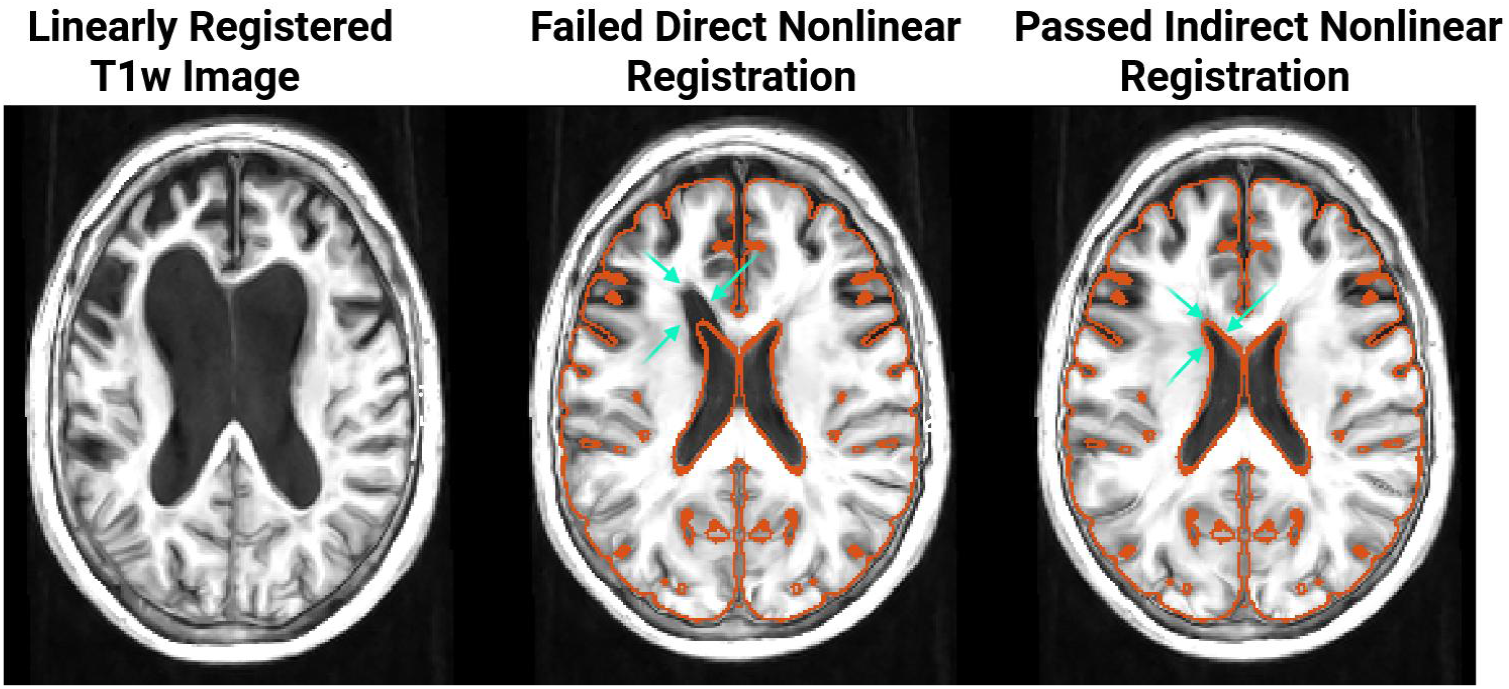
Example of a NIFD participant with frontotemporal dementia and severe atrophy levels that failed direct nonlinear registration to the MNI-ICBM152 average template, but passed indirect nonlinear registration using MNI-FTD136^25^ average template as an intermediate registration target. Cyan arrows show the area where direct nonlinear registration failed to capture the severity of ventricular expansion but indirect registration was successful.

Table 3 summarizes the results of failure rate comparisons between the *PELICAN* and *FreeSurfer. FreeSurfer* had consistently higher failure rates across all datasets for both linear registration and tissue segmentation, with values ranging between 1.55% to 24.4% versus 0.001% to 3.75% for linear registration (all p-values < 0.005), and 22.73% to 34.42% versus 0.83% to 3.77% for tissue segmentation (all p-values < 0.001).

**Table 3.** PELICAN and FreeSurfer failure rates for a subset of the datasets included in this study. Failure rates are reported in percentages of all cases.

| Dataset | UK BioBank | NIFD | ADNI | SIMON |
| --- | --- | --- | --- | --- |
| N | 244 | 751 | 232 | 53 |
| Linear registration - PELICAN | 3.75 | 0.001 | 2.16 | 1.89 |
| Linear registration - FreeSurfer | 24.4 | 1.55 | 19.82 | 12.24 |
| Tissue Segmentation - PELICAN | 0.83 | 1.59 | 1.72 | 3.77 |
| Tissue Segmentation - FreeSurfer | 34.42 | 22.73 | 23.27 | 29.18 |

Table 4 summarizes the results of the reliability comparisons between the *PELICAN* and *FreeSurfer* for SIMON dataset, reflecting the amount of variability in volumetric estimations of the two pipelines for the cases that had passed all QC steps for both pipelines. Given the nature of the dataset (i.e. all images were acquired within a year), in an ideal case, the variability in measurements should be minimal. *PELICAN* outperformed FreeSurfer, showing smaller amounts of variability in eTIV (1.1% for *PELICAN* versus 3.0% for *FreeSurfer*) as well as volumetric estimates in native space (i.e. not impacted by eTIV variability). Variability values in tissue volumes were below 2.5% for *PELICAN* and below 3.8% for FreeSurfer, except for WMHs (15% for PELICAN versus 25.6% for FreeSurfer), for which the higher variability rates were likely due to the very low overall WMH burden. Of note, the eTIV was very different between the two methods. PELICAN-estimated mean eTIV was 1,382,794, which was very close to the sum of all tissue types estimated by PELICAN (1,380,031, i.e. 0.002% difference), whereas FreeSurfer-estimated eTIV (1,527,046) was significantly larger than the sum of all tissue types estimated by FreeSurfer (1,218,104, i.e. 25.36% difference). Figure 4 shows an example of PELICAN and FreeSurfer segmentations as well as their differences for a SIMON image.

**Table 4.** PELICAN and FreeSurfer reliability comparisons for the SIMON dataset. Results are reported as raw values as well as standard deviations normalized over mean volumes.

| Dataset | PELICAN | FreeSurfer |
| --- | --- | --- |
| eTIV | 1,382,794 ± 15,919.6 (0.011) | 1,527,046 ± 46,076 (0.030) |
| Cortical Gray Matter | 676,203.7 ± 14,498.2 (0.021) | 516,324.1 ± 11,449.0 (0.022) |
| Deep Gray Matter | 47,449.5 ± 414.4 (0.008) | 50,998.9 ± 1,264.1 (0.025) |
| White Matter | 400,026.3 ± 5,132.7 (0.012) | 456,122.8 ± 6,579.4 (0.015) |
| Ventricle | 10,592.5 ± 260.9 (0.025) | 23,433.2 ± 624.4 (0.027) |
| WMH | $601.1 \pm 90.2$ (0.150) | $1,052.9 \pm 269.6$ (0.256) |
| Cerebellar Gray Matter | $115,942.1 \pm 2,161$ (0.018) | $111,195.5 \pm 3,082.6$ (0.028) |
| Cerebellar White Matter | $25,330.9 \pm 349.1$ (0.013) | $28,983.1 \pm 1,092.2$ (0.038) |

**Figure 4.**
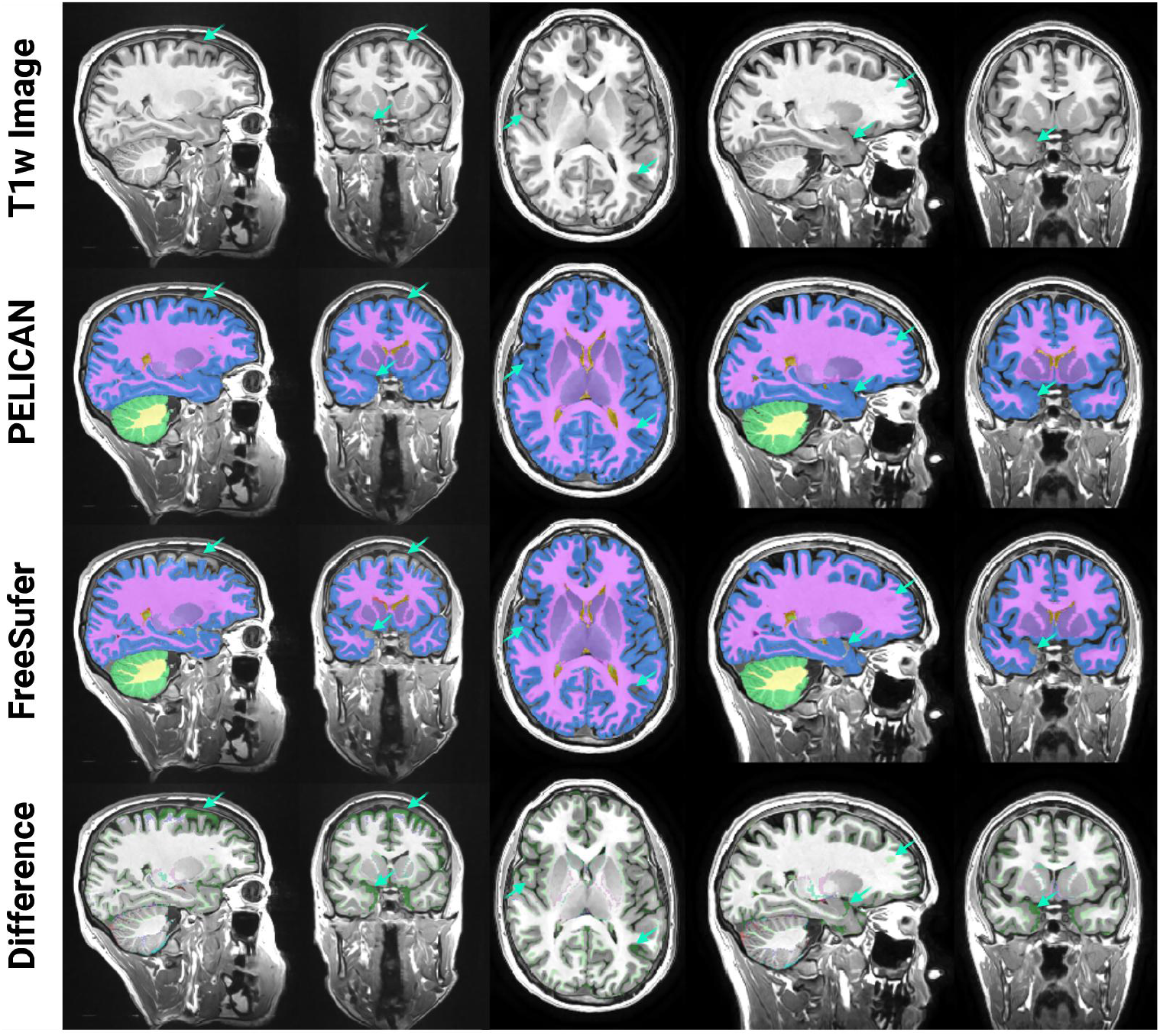
Comparison of PELICAN versus FreeSurfer segmentations for a SIMON image. **First row:** axial, sagittal, and coronal slices of the preprocessed T1w images. **Second row:** PELICAN segmentation overlaid on the T1w image. **Third row:** FreeSurfer segmentation overlaid on the T1w image. **Fourth row:** areas of difference between PELICAN and FreeSufer segmentations. **Cyan arrows** show main areas of difference between the two segmentations, mainly reflecting the tendency of FreeSurfer to undersegment cortical gray matter, whereas PELICAN favors oversegmentation.

## Discussion

In this work, we present PELICAN, our robust multi-modal longitudinal image processing pipeline. We validated the performance of PELICAN in over 35,000 scans from over 24,000 healthy aging individuals as well as patients with different neurodegenerative disorders based on visual quality assessments. We further compared the performance and reliability of the PELICAN against the registrations and volumetric measurements of FreeSurfer, demonstrating lower failure rates and greater reliability for the PELICAN.

Each step of PELICAN was designed based on our prior experience with longitudinal image processing of multi-center and multi-scanner MRI data from aging and neurodegenerative disease populations. Different components and steps of the pipeline were further refined based on our extensive visual quality assessments. For example, while it is not a requirement of the specific tools, we perform both intensity non-uniformity correction and intensity normalization using an initial brain mask that is derived based on an initial set of linear registrations. While computationally more expensive, this decision ensures that signals from non-brain tissues (e.g. scalp or neck fat, vitamin pills used to identify laterality, etc.) or potential presence of artifacts outside of the brain do not impact the outputs of these normalization steps. This step likely contributes to the high quality of the derived brain masks and segmentations.

We combined several MINC-based tools with linear and nonlinear registration tools from *ANTs* to create *PELICAN*. This decision was based on extensive comparisons and visual quality control of MINC-based and ANTs-based linear and nonlinear registration tools. While some variants of the linear registration methods provided by the MINC toolkit have excellent performance overall^9^, they were outperformed by ANTs linear registration in cases with poor initial head positioning, for which the pipeline required to estimate large rotation parameters to perform accurate registrations. As this was a common occurrence in datasets such as the UK BioBank, we opted to provide the default version of the pipeline with ANTs linear registration, while we also provide a second open source version using a MINC-based linear registration technique in the same repository. With regards to nonlinear registration, it is widely acknowledged that the ANTs implementation can provide robust and accurate results in a wide range of applications, and was thus used in all our nonlinear registration steps.

While there are also a number of other open access DBM and VBM pipelines available, we chose to compare the performance of *PELICAN* against *FreeSurfer* for several reasons: First, *FreeSurfer* is the most widely used pipeline for deriving volumetric and regional brain measurements, including in large-scale consortium efforts such as the ENIGMA studies^40–43^. Second, *FreeSurfer* also enables longitudinal analyses which was a particular point of interest. Third, an important concern when performing T1w-based image segmentations in age and neurodegenerative disease populations is presence of WMHs, which can result in erroneous tissue segmentations^17^. Our T1w-based tissue segmentations are derived based on *BISON*, which segments WMHs as a separate class, and is thus able to provide accurate tissue segmentations even in cases with extensive WMH pathology^18^. This is also the case for *FreeSurfer* which labels WMHs as a separate class named T1w hypointensities, and is thus likely more accurate compared to methods that do not label WMHs as a separate class^18^. As such, we believe that it would be more relevant to compare *PELICAN* segmentations against *FreeSurfer*. Finally, we were interested in pipelines that can be deployed on high performance computing clusters and enable parallel analyses of large scale datasets such as the UK BioBank. This requirement in particular made MATLAB-based pipelines such as *SPM* and *CAT12* prohibitive as they required MATLAB licenses. As such, we opted to not include *CAT12* in our comparisons. Nevertheless, we processed the NIFD dataset with *CAT12* on a local computer and examined the outputs, finding substantial inaccuracies in the results (Supplementary Figure S.12).

To assess the robustness of PELICAN’s performance, we applied it to data from several different aging and neurodegenerative disease cohorts^44–53^, and performed extensive visual quality controls on all outputs. While we have reported PELICAN’s failure rates in the resulting articles and have generally found similar performance levels across all datasets, we present the results from 5 specific datasets here: i) the ADNI, as a widely used multi-site and multi-scanner Alzheimer’s disease dataset with a harmonized protocol ii) the UK Biobank, as a widely used aging dataset with high proportions of challenging cases for linear registration (due to poor initial head positioning), iii) NIFD, as an open access dataset including many cases with severe atrophy that are usually challenging for nonlinear registration, iv) NACC, as another open access dataset including images from a wide range of scanner manufacturers, models, imaging protocols, and resolutions (i.e. no MRI protocol harmonization), and v) SIMON, to assess measurement reliability.

*PELICAN* consistently outperformed *FreeSurfer* in linear registrations, with failure rates ranging between 0.001% to 3.75%, compared to 1.55%-24.4% (Table 3). This is particularly important as both *FreeSurfer*^54,55^ (https://surfer.nmr.mgh.harvard.edu/fswiki/eTIV) and *PELICAN* estimate total intracranial volume based on linear registrations, and any errors in linear registration would result in errors in the eTIV estimates. Based on previous reports by Klasson and colleagues^54^ as well as others^56,57^, inaccuracies in FreeSurfer eTIV estimates can also be systematic, as evidenced by positive correlations between eTIV and total brain volume when controlling for intracranial volume calculated by delineating the dura mater. Since eTIV estimates are commonly used to account for head size differences^58–60^, it is crucial to quality control the linear registrations to ensure their accuracy. Furthermore, even in the subset of the SIMON images that passed quality control for both pipelines, *FreeSurfer* eTIV estimations showed lower reliability compared to *PELICAN* (Table 4).

*PELICAN* also outperformed *FreeSurfer* in tissue segmentation, with consistently lower failure rates. While *FreeSurfer* had many drastically failed cases (e.g. Supplementary Figure S.10), *PELICAN* failures were more subtle (e.g. Supplementary Figure S.9). *PELICAN* incorporates tissue segmentation based on *BISON*^18^, which receives preprocessed images, brain masks, and linear + nonlinear registration transformations as input, and assigns tissue labels ranging between 1 (CSF) to 9 (WMH) to all voxels inside the brain mask. As such, voxels that are not included in the brain mask will not be segmented by *BISON*. Similarly, if non-brain regions (e.g. skull and dura) are included in the brain mask, they will inevitably be segmented incorrectly (usually as CSF or cortical GM). This is generally uncommon as *BEaST* failure rates are usually low^38^ (Table 2). Nevertheless, all outputs should be carefully quality controlled for any pipeline prior to use. Furthermore, *BISON* uses the registration transformations to calculate spatial prior features, and as such, while small inaccuracies are usually well tolerated, gross failures in linear and nonlinear registrations would result in segmentation failures.

We did not include *FreeSurfer* brain masks in our comparisons as *FreeSurfer* brain extractions are particularly error prone (e.g. Supplementary Figure S.11)^54,61,62^, and would have resulted in very high failure rates. Furthermore, *FreeSurfer* segmentations and cortical thickness estimations do not directly rely on *FreeSurfer* brain masks, and as such, the poor quality of the *FreeSurfer* brain masks does not result in failure in consequent steps and would not be a fair representation of the overall performance of the pipeline. Nevertheless, we do not recommend use of *FreeSurfer* brain masks in any downstream analyses. It is also important to note that for *PELICAN* and other similar DBM and VBM pipelines, accuracy of the brain masks is important as it can translate into inaccuracies in the estimated nonlinear transformations, since brain masks are used as inputs for the nonlinear registration. As such, careful quality control of the brain masks is warranted to ensure accuracy of the DBM and VBM estimations.

Finally, *PELICAN* also consistently outperformed *FreeSurfer* with respect to reliability, as measured using SIMON dataset, showing lower amounts of variability in eTIV (0.011 vs 0.030) as well as gray matter, white matter, and WMH volumes. Note that the CSF was not included in this comparison, as *FreeSurfer* does not consistently segment the CSF. Overall, *FreeSurfer* provides significantly larger eTIV estimates compared to *PELICAN*, whose estimates were closer to the sum of its estimated tissue volumes (as well as *FreeSurfer*’s). This might result from a difference in what *FreeSurfer* and *PELICAN* consider the intracranial space, or the fact that the *FreeSurfer* estimations are made based on the MNI-305 template (https://surfer.nmr.mgh.harvard.edu/fswiki/eTIV) which is larger than the MNI-ICBM152. Regardless, as long as eTIVs are derived using the same software, this normalization should not impact the results as the two measures are highly correlated (e.g. r = 0.94, p < 0.0001 in the UK Biobank dataset). However, *FreeSurfer*-based eTIV estimates should not be compared against or combined with eTIV estimates from other softwares. Furthermore, in partial volume areas such as narrow sulci, *FreeSurfer* tended to favour undersegmenting the cortical gray matter, whereas *PELICAN* favoured oversegmenting the cortical gray matter and undersegmenting the CSF, resulting in overall larger estimated gray matter volumes by *PELICAN* compared to *FreeSurfer*.

One of the novel features implemented in *PELICAN* is its capability to use population specific average templates as intermediate targets for the nonlinear registration step. This step can significantly improve nonlinear registrations for cases with substantial levels of atrophy (Figure 3 as well as Supplementary Figure S.13), allowing for their inclusion in downstream analyses. This is particularly important when studying neurodegenerative disorders such as frontotemporal dementia, as exclusion of cases with extensive levels of atrophy due to their higher failure rates would inevitably lead to underestimation of the atrophy levels in these populations. Our results showed lower failure rates of indirect compared to direct nonlinear registrations for NIFD (1.83% versus 6.34%) and ADNI (1.86% versus 5.88%), half of which included individuals with frontotemporal dementia and Alzheimer’s dementia, respectively. We have made both MNI-FTD136 and ADNI templates available as intermediate registration targets with PELICAN, since based on our experience, severity of atrophy in individuals with frontotemporal dementia and Alzheimer’s dementia commonly necessitates use of intermediate templates to improve the nonlinear registrations. However, *PELICAN* also accepts other population specific average templates as the intermediate targets.

Accurate and robust longitudinal image processing is essential for many neuroimaging applications, and can improve sensitivity to detection of the more subtle brain changes that might not be detected using pipelines that are cross-sectional or not suited to the populations of interest. Our results suggest that *PELICAN* can provide accurate and robust registration, segmentation, and morphometry maps that can be particularly useful when studying aging and neurodegenerative disease populations.

## Acknowledgements

The authors would like to thank Dr. Gabriel Devenyi for creating the singularity container image of ANTs, MINC, and Anaconda tools, that is used to run PELICAN as well as for his valuable discussions and feedback. The authors also acknowledge Digital Research Alliance of Canada (https://www.alliancecan.ca/en) for the usage of the computing resources in the current work. Data collection and sharing for this project was funded by the Alzheimer Disease Neuroimaging Initiative (ADNI) (NIH grant U01 AG024904) and DOD ADNI (Department of Defense award number W81XWH-12-2-0012). The ADNI is funded by the National Institute on Aging, the National Institute of Biomedical Imaging and Bioengineering, and through generous contributions from the following: AbbVie, Alzheimer Association; Alzheimer Drug Discovery Foundation; Araclon Biotech; BioClinica, Inc.; Biogen; Bristol-Myers Squibb Company; CereSpir, Inc.; Cogstate; Eisai Inc.; Elan Pharmaceuticals, Inc.; Eli Lilly and Company; EuroImmun; F. Hoffmann-La Roche Ltd. and its affiliated company Genentech, Inc.; Fujirebio; GE Healthcare; IXICO Ltd.; Janssen Alzheimer Immunotherapy Research & Development, LLC; Johnson & Johnson Pharmaceutical Research & Development LLC; Lumosity; Lundbeck; Merck & Co., Inc.; Meso Scale Diagnostics, LLC; NeuroRx Research; Neurotrack Technologies; Novartis Pharmaceuticals Corporation; Pfizer Inc.; Piramal Imaging; Servier; Takeda Pharmaceutical Company; and Transition Therapeutics. The Canadian Institutes of Health Research is providing funds to support ADNI clinical sites in Canada. Private sector contributions are facilitated by the Foundation for the National Institutes of Health (fnih.org). The grantee organization is the Northern California Institute for Research and Education, and the study is coordinated by the Alzheimer Therapeutic Research Institute at the University of Southern California. ADNI data are disseminated by the Laboratory for Neuro Imaging at the University of Southern California. The NACC database is funded by NIA/NIH Grant U24 AG072122. NACC data are contributed by the NIAfunded ADRCs: P30 AG062429 (PI James Brewer, MD, PhD), P30 AG066468 (PI Oscar Lopez, MD), P30 AG062421 (PI Bradley Hyman, MD, PhD), P30 AG066509 (PI Thomas Grabowski, MD), P30 AG066514 (PI Mary Sano, PhD), P30 AG066530 (PI Helena Chui, MD), P30 AG066507 (PI Marilyn Albert, PhD), P30 AG066444 (PI David Holtzman, MD), P30 AG066518 (PI Lisa Silbert, MD, MCR), P30 AG066512 (PI Thomas Wisniewski, MD), P30 AG066462 (PI Scott Small, MD), P30 AG072979 (PI David Wolk, MD), P30 AG072972 (PI Charles DeCarli, MD), P30 AG072976 (PI Andrew Saykin, PsyD), P30 AG072975 (PI Julie A. Schneider, MD, MS), P30 AG072978 (PI Ann McKee, MD), P30 AG072977 (PI Robert Vassar, PhD), P30 AG066519 (PI Frank LaFerla, PhD), P30 AG062677 (PI Ronald Petersen, MD, PhD), P30 AG079280 (PI Jessica Langbaum, PhD), P30 AG062422 (PI Gil Rabinovici, MD), P30 AG066511 (PI Allan Levey, MD, PhD), P30 AG072946 (PI Linda Van Eldik, PhD), P30 AG062715 (PI Sanjay Asthana, MD, FRCP), P30 AG072973 (PI Russell Swerdlow, MD), P30 AG066506 (PI Glenn Smith, PhD, ABPP), P30 AG066508 (PI Stephen Strittmatter, MD, PhD), P30 AG066515 (PI Victor Henderson, MD, MS), P30 AG072947 (PI Suzanne Craft, PhD), P30 AG072931 (PI Henry Paulson, MD, PhD), P30 AG066546 (PI Sudha Seshadri, MD), P30 AG086401 (PI Erik Roberson, MD, PhD), P30 AG086404 (PI Gary Rosenberg, MD), P20 AG068082 (PI Angela Jefferson, PhD), P30 AG072958 (PI Heather Whitson, MD), P30 AG072959 (PI James Leverenz, MD). Data collection and sharing for this project was also funded by the Frontotemporal Lobar Degeneration Neuroimaging Initiative (National Institutes of Health Grant R01 AG032306). The study is coordinated through the University of California, San Francisco, Memory and Aging Center. FTLDNI data are disseminated by the Laboratory for Neuro Imaging at the University of Southern California. This research was conducted using the UKBB Resource under approved application 45551. We thank the UKBB participants and team for their work in collecting, processing, and disseminating these data for analysis.

