## Supplementary Figures for "PELICAN: a Longitudinal Image Processing Pipeline for Analyzing Structural Magnetic Resonance Images in Aging and Neurodegenerative Disease Populations"

### Supplementary Materials

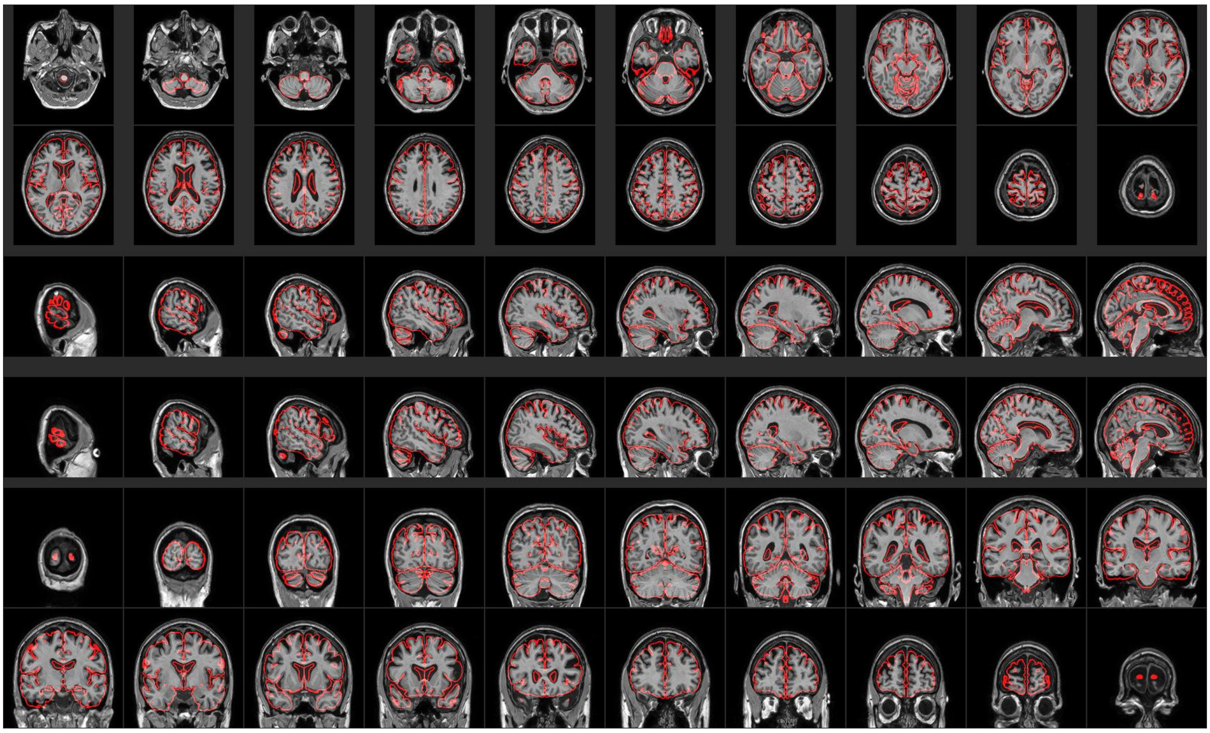

Figure S.1. Example of a passed quality control image for linear registration of a T1w image.

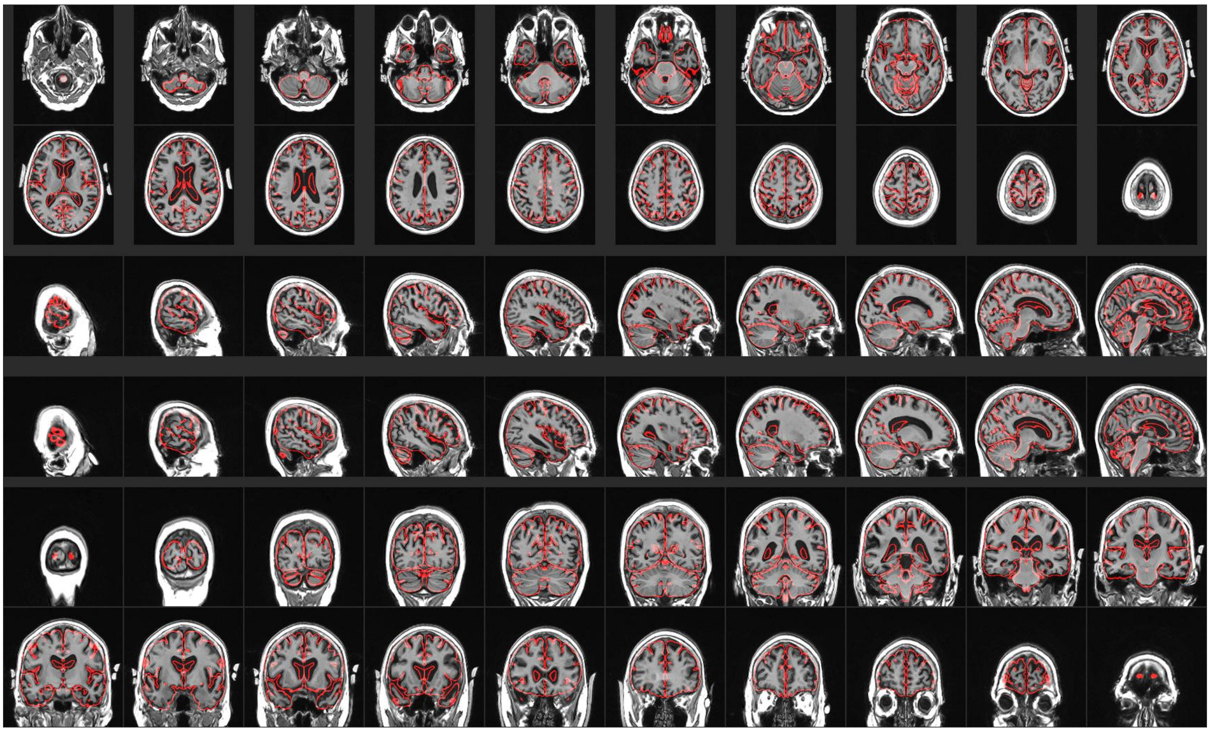

Figure S.2. Example of a failed quality control image for linear registration of a T1w image.

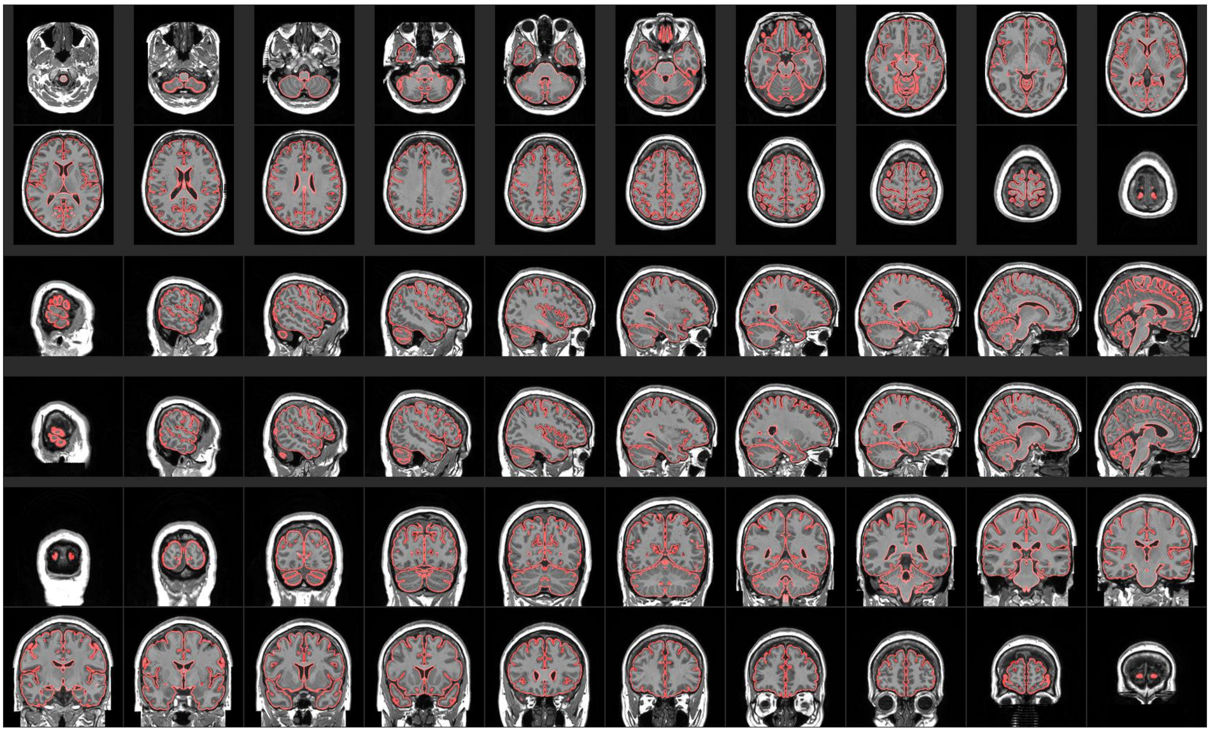

Figure S.3. Example of a passed quality control image for direct nonlinear registration of a T1w image.

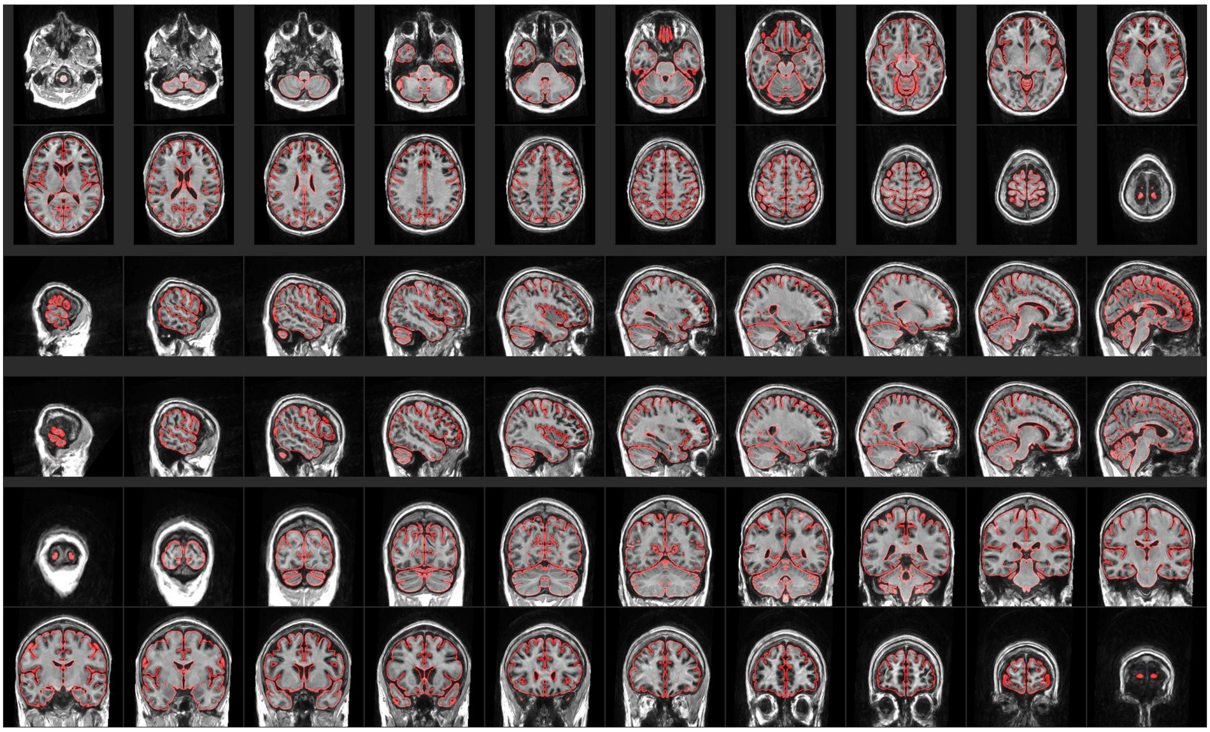

Figure S.4. Example of a failed quality control image for direct nonlinear registration of a T1w image.

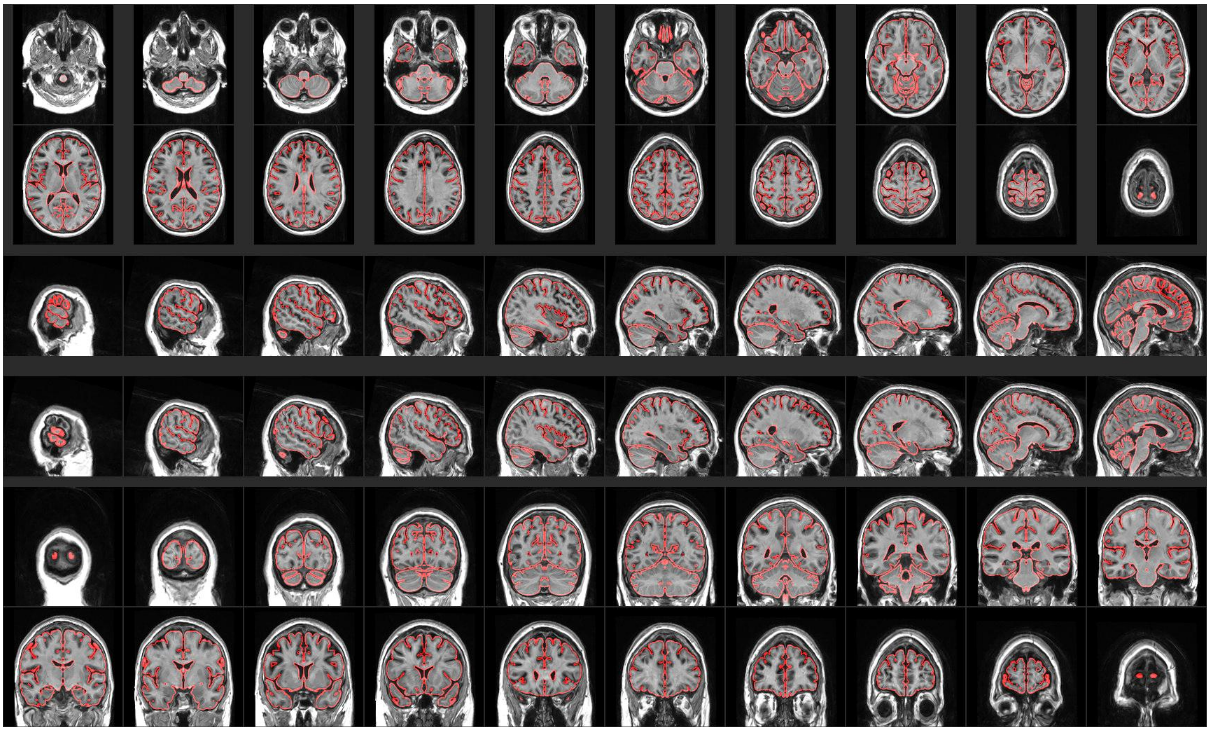

Figure S.5. Example of passed indirect nonlinear registration of the same T1w image presented in Figure S.10.

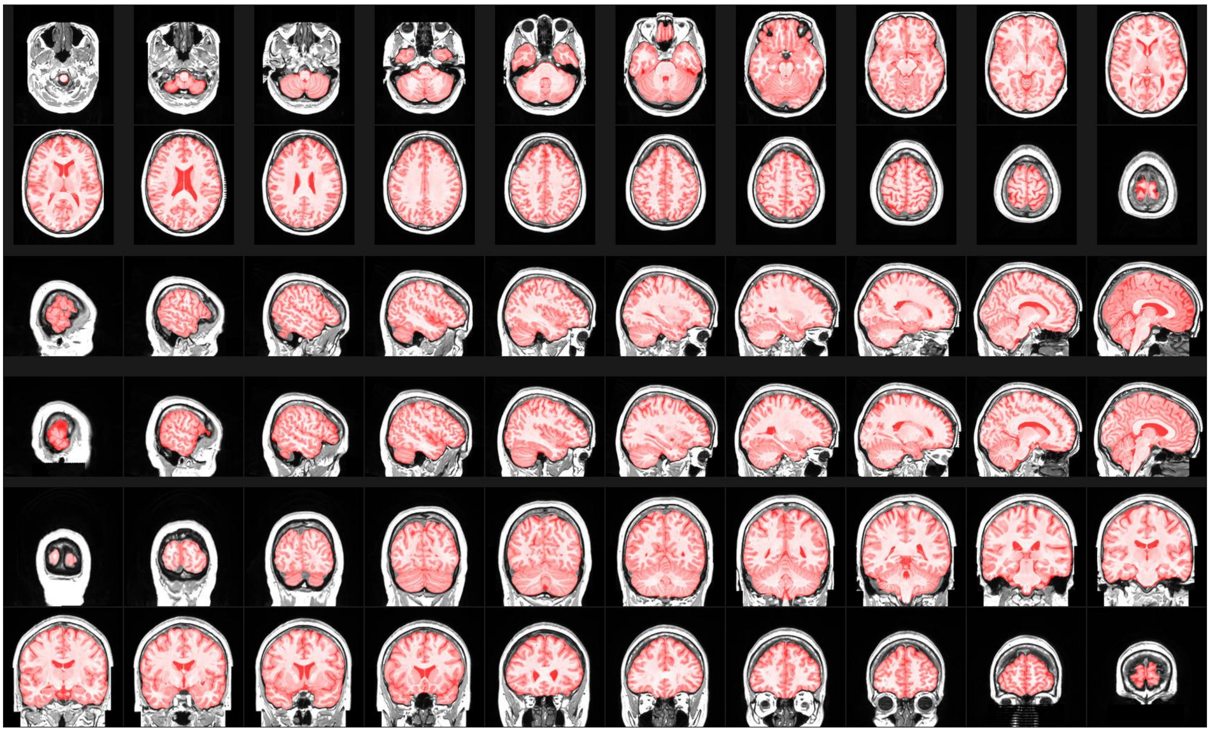

Figure S.6. Example of a passed quality control image for BEaST brain mask overlaid on the linearly registered T1w image.

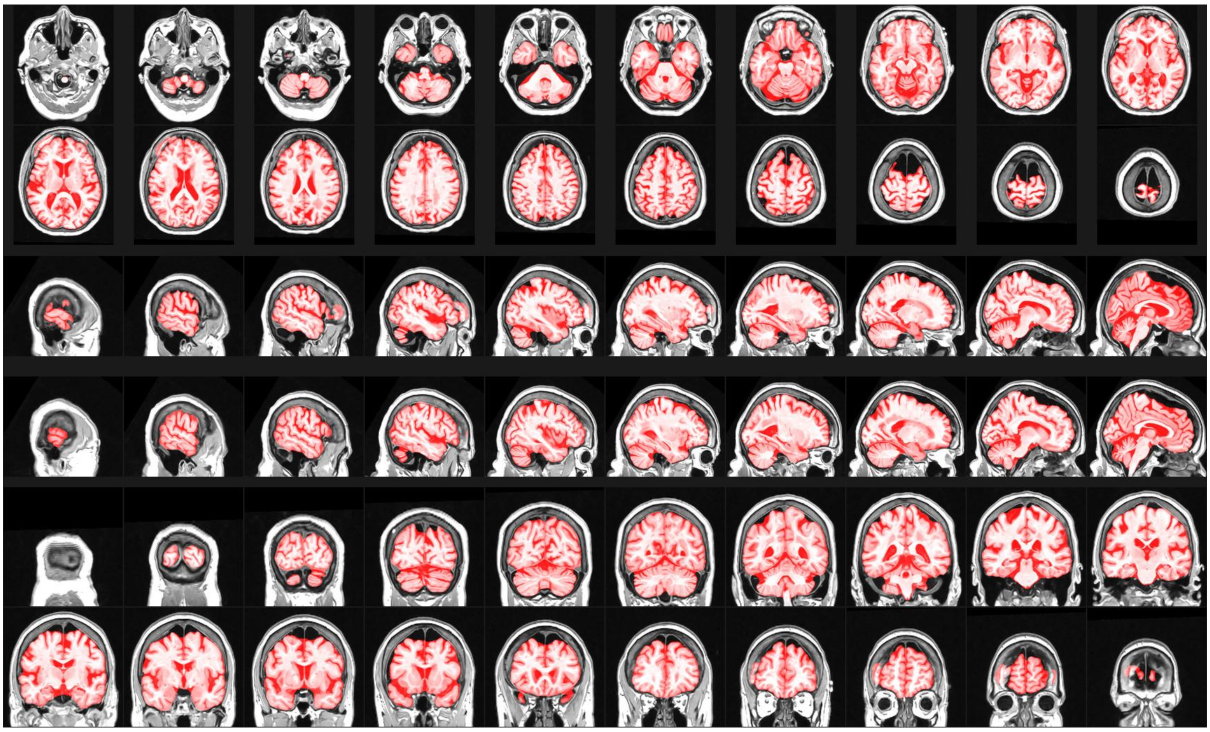

Figure S.7. Example of a failed quality control image for BEaST brain mask overlaid on the linearly registered T1w image.

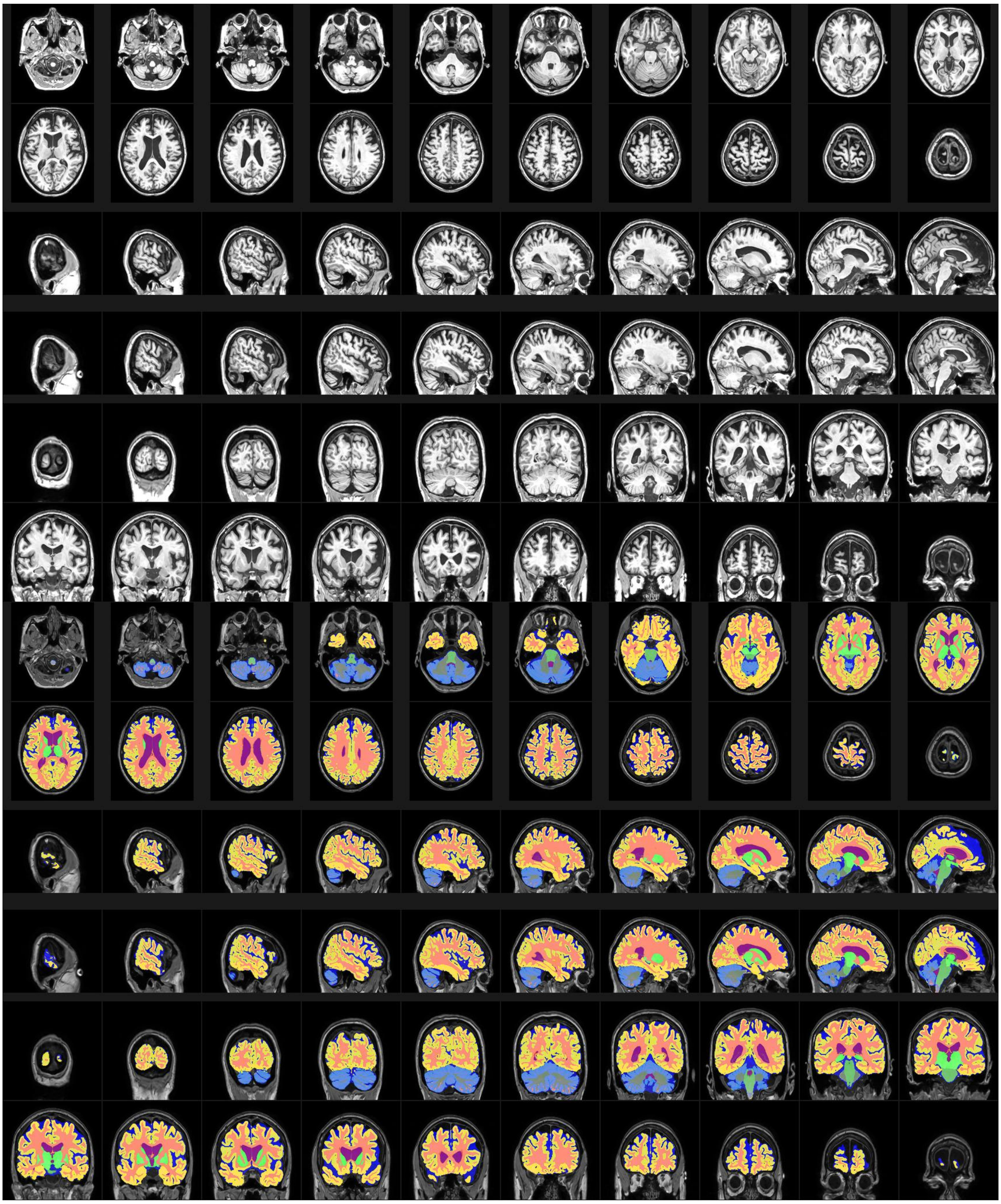

Figure S.8. Example of a passed quality control image for BISON mask overlaid on the linearly registered T1w image.

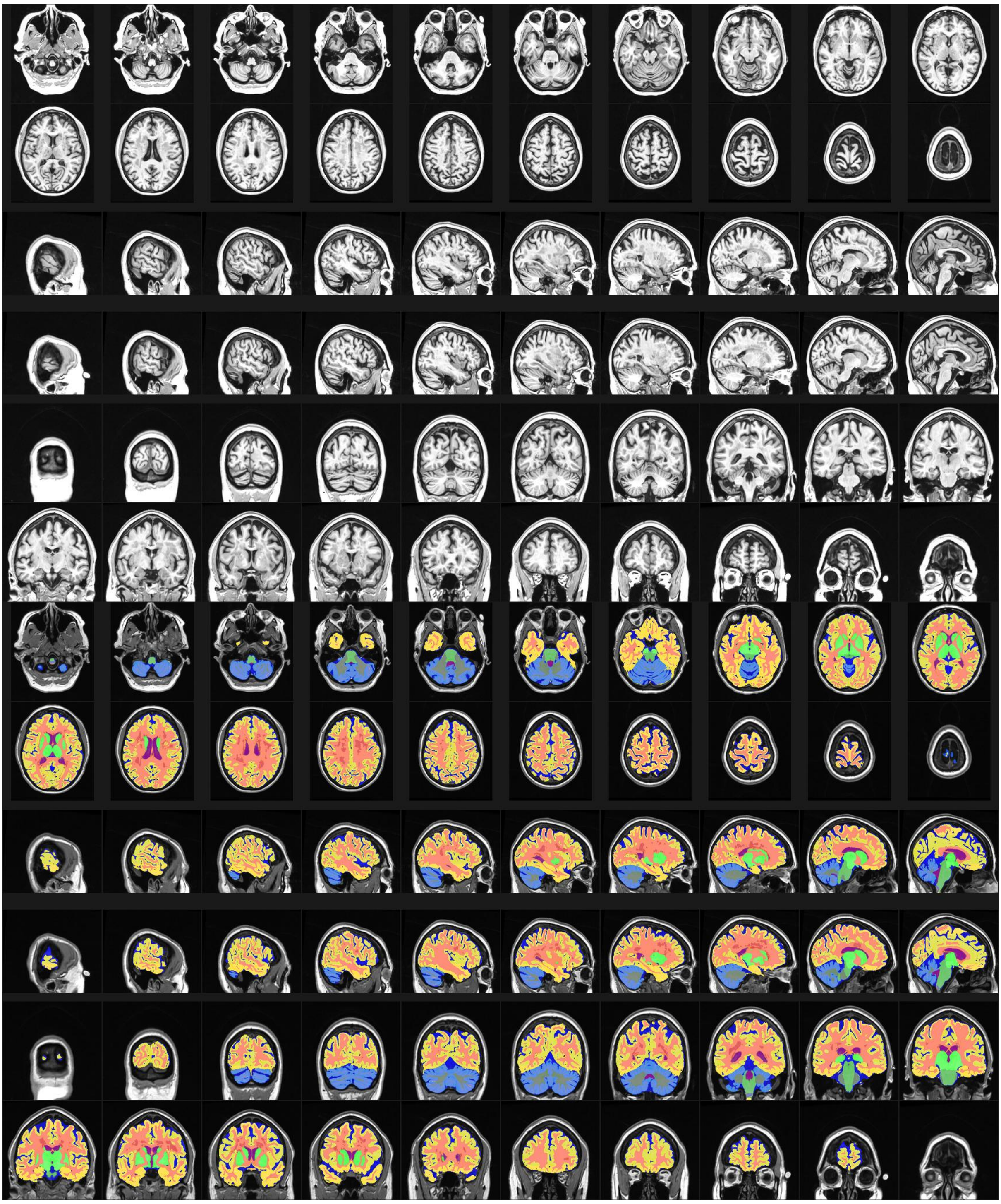

Figure S.9. Example of a failed quality control image for BISON  mask overlaid on the linearly registered T1w image.

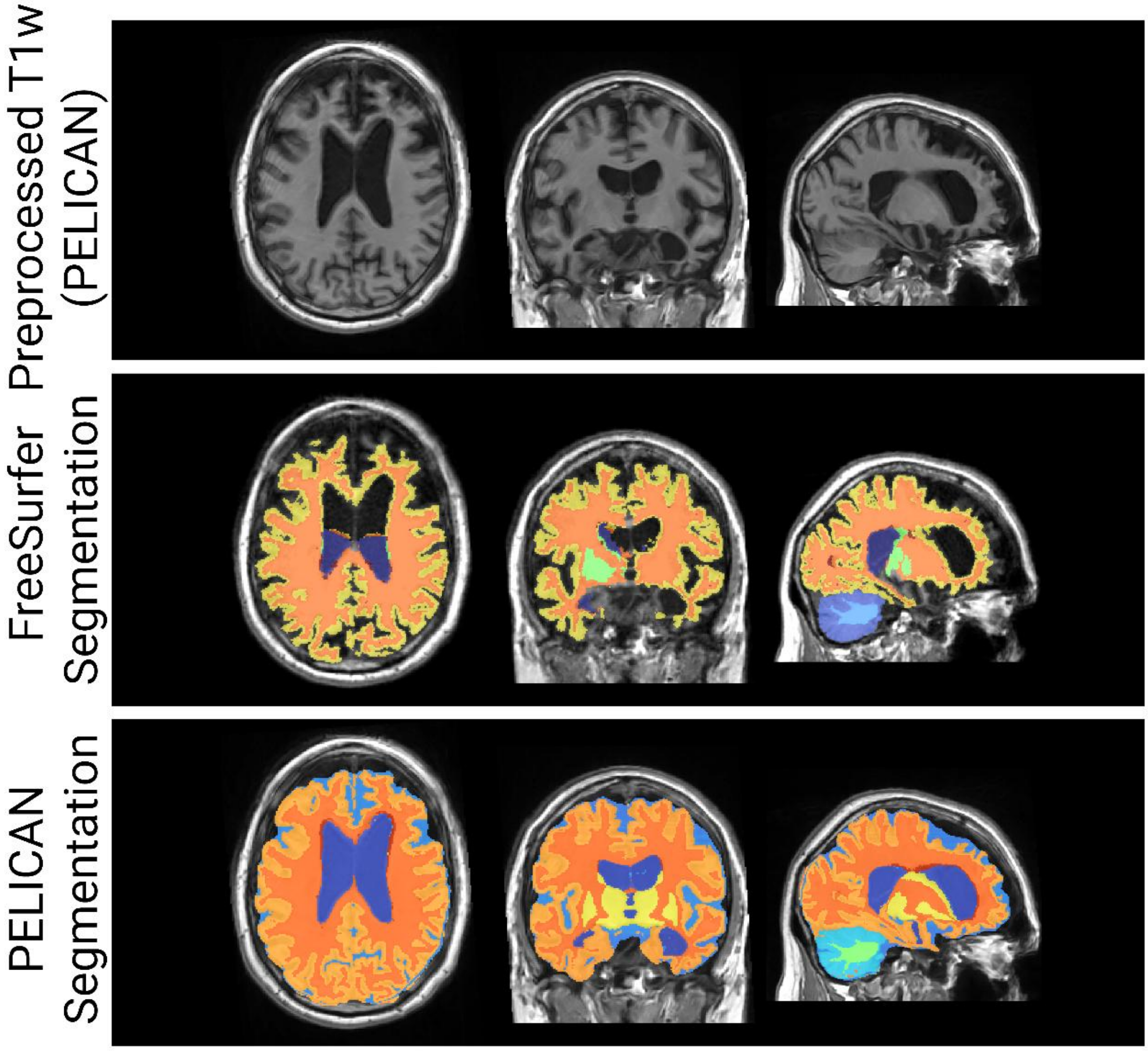


Figure S.10. Example of a failed quality control image for FreeSurfer tissue segmentations and corresponding passed PELICAN segmentations overlaid on the linearly registered T1w image of an individual with frontotemporal dementia from the NIFD dataset. Major errors in FreeSurfer segmentations included not fully capturing the frontal and temporal lobes as well as segmenting periventricular WMHs as cortical gray matter.

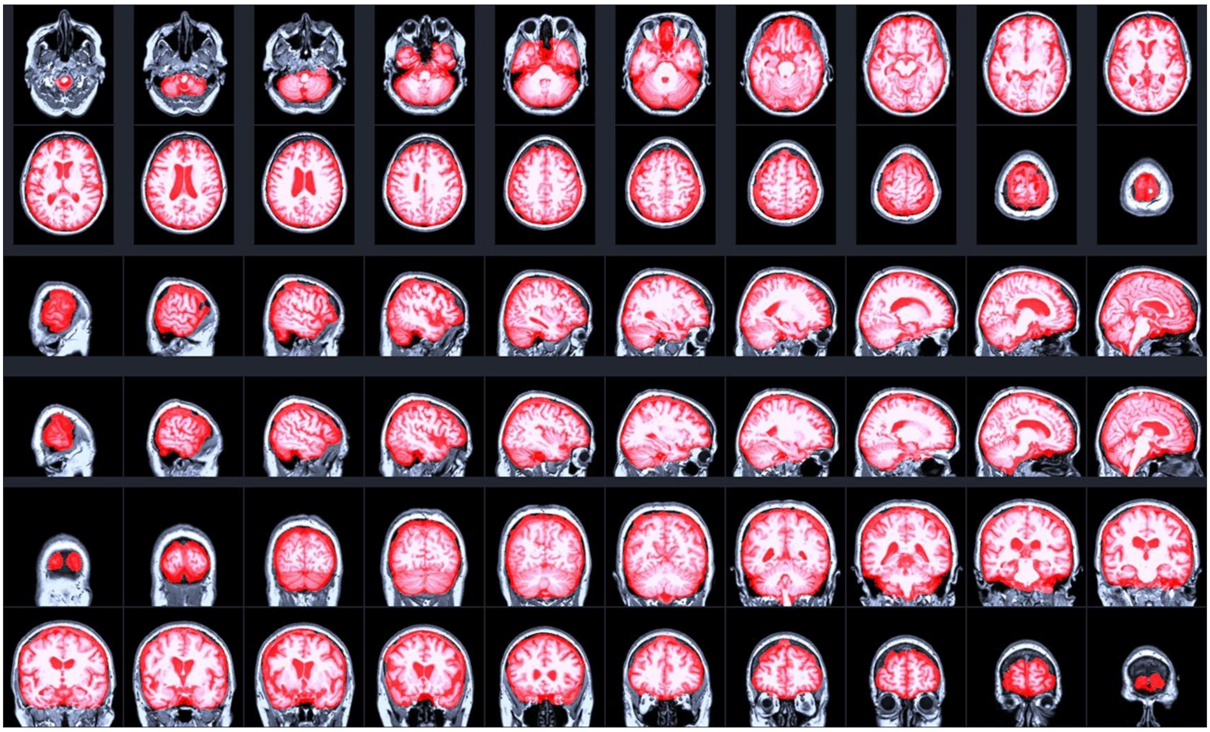

Figure S.11. Example of a failed quality control image for FreeSurfer brain mask overlaid on the linearly registered T1w image. Note the inclusion of non brain tissues in the mask.

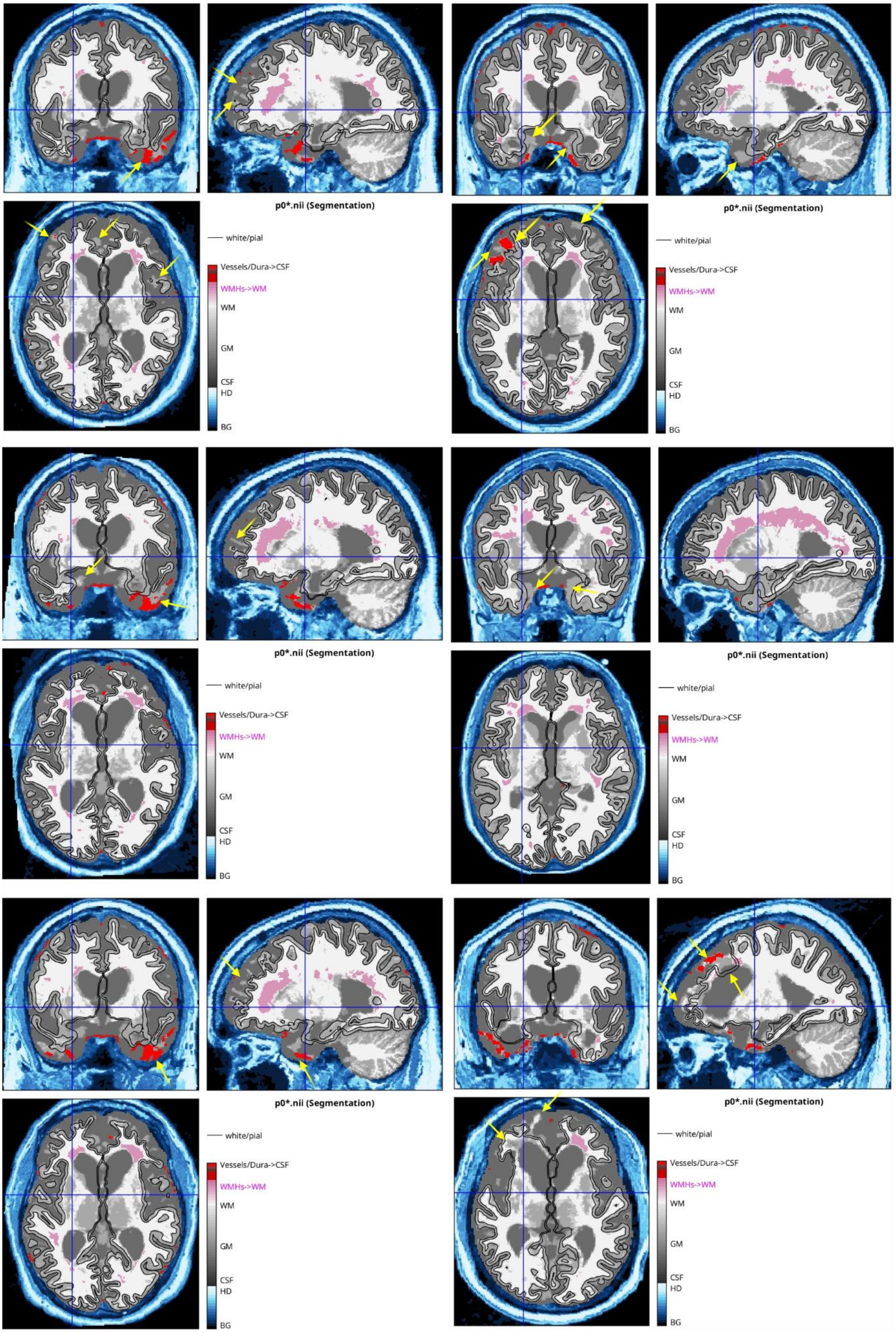


Figure S.12. Example of a failed quality control image for FreeSurfer tissue segmentation mask overlaid on the linearly registered T1w image. Major failures include not capturing frontal and temporal lobe cortical regions in areas of severe atrophy, as indicated by yellow arrows.

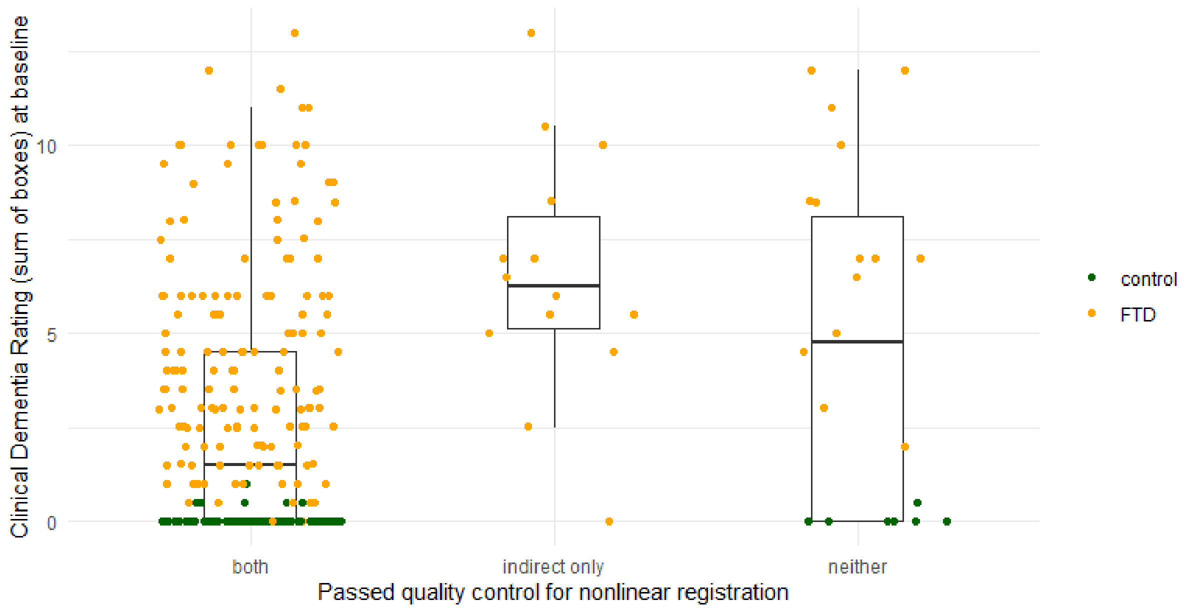


Figure S.13. Clinical dementia rating sum of boxes (CDR-SB) values for cases that passed both direct and indirect nonlinear registration (left), cases that only passed indirect nonlinear registration (middle), and cases that failed both registrations (right) for the NIFD dataset. CDR-SB values reflect disease severity, suggesting that the cases that failed direct but passed indirect registration were patients with greater levels of disease severity. FTD: frontotemporal dementia.
